# PET-Microplastics Trigger Endothelial Glycocalyx Loss via ER Stress and ROS Unleashing IL-1β-Driven SMC Switching and Early Aortic Structural Impairment

**DOI:** 10.1101/2025.09.22.677783

**Authors:** Weixue Huo, Jin Qu, Sen Wang, Mengwei He, Zhaoxiang Zeng, Deping Kong, Lushun Yuan, Rui Feng

## Abstract

Polyethylene terephthalate microplastics (PET-MPs), a major microplastics component identified in human vasculature, pose emerging environmental health risks. This study systemically profiled MPs in human aortic tissues and investigated the mechanisms underlying PET-MPs-induced aortic injury in vivo and in vitro. Chronic oral exposure of Sprague-Dawley rats to PET-MPs (1.0-100 mg/L) resulted in endothelial glycocalyx loss and structural impairment of aortic elastic fibers, with MPs accumulating within aortic endothelial cells. Transcriptomic and biochemical analyses revealed that PET-MPs triggered endoplasmic reticulum stress (ERS) and reactive oxygen species (ROS) generation in human aortic endothelial cells (HAECs), driving glycocalyx degradation and NF-κB-mediated inflammation. Proteomic profiling identified endothelial-derived IL-1β as a key mediator, which subsequently induced phenotypic switching in human aortic smooth muscle cells (HASMCs) in vitro. Pharmacological inhibition of ERS (TUDCA), ROS (NAC), or IL-1β (Canakinumab) attenuated this pathogenic cascade. Crucially, restoration of the glycocalyx using Sulodexide mitigated endothelial dysfunction and downstream HASMC phenotypic switching. These findings establish endothelial glycocalyx degradation via ERS-ROS as a novel mechanism for PET-MPs-induced vascular injury and highlight glycocalyx protection as a potential strategy against environmental microplastic hazards.

## 1. Introduction

Microplastics (MPs) are pervasive environmental hazards, recently reported to contaminate human tissues. MP components have been detected in human blood, placenta, liver, kidneys, and blood vessels(1–3), with estimated daily intake reaching 0.23-11.9 mg/kg.(4–6) Multiple studies indicate that MPs pose potential toxic hazards to the digestive, respiratory, and immune systems. Critically, emerging evidence demonstrated that MPs can cause vascular damage(7–10), yet the key microplastic components involved and their specific effects on aortic integrity are not yet characterized.

All vascular lumens, including the aorta, are lined with a glycoprotein and polysaccharide complex termed the endothelial glycocalyx.(11) The endothelial glycocalyx is a bush-like structure lining the vascular lumen in a gel-like state, separating blood from tissues. It serves multiple functions, including regulating vascular permeability, maintaining smooth blood flow, transducing mechanical forces within vessels, and inhibiting blood coagulation and leukocyte adhesion.(11–16) Notably, the endothelial glycocalyx is a labile and fragile structure susceptible to disruption by various stimuli. Sepsis, ischemia-reperfusion injury, trauma, diabetic nephropathy, and hyperlipidemia can all cause glycocalyx damage, subsequently leading to vascular injury and pathogenesis.(17–24)

Endoplasmic reticulum stress (ERS) represents a critical pathway for environmental toxin-induced vascular injury. As the primary site for protein folding in endothelial cells, ER dysfunction upon toxicant exposure triggers the unfolded protein response (UPR).(25, 26) Seminal work by Kaufman *et al.* established that UPR activation elevates reactive oxygen species (ROS) generation, initiating a self-amplifying ERS-ROS cascade that disrupts vascular homeostasis.(27, 28) Importantly, this axis directly compromises endothelial barrier integrity by degrading the glycocalyx. Elevated ROS activating matrix metalloproteinases (MMPs) accelerates glycocalyx shedding.(29, 30) This dual assault erodes endothelial junctional stability and increases permeability-key initiators of aortic structural failure. While ERS is known to drive endothelial activation in sepsis and diabetes (31, 32), its role in microplastic-induced vascular injury and the aortic extracellular matrix degradation remains uncharacterized.

The specific composition of microplastics is complex. The main types of current MP pollutants include polystyrene (PS), polyethylene terephthalate (PET), polyethylene (PE), polyvinyl chloride (PVC), among others.(3, 10, 33–35) Interestingly, we note significant differences in the abundance of various MP types across different human tissues and body fluids. The content and composition of MPs differ in the testes, semen, endometrium, bone, muscle tissue, and blood.(36–39) The primary components of MPs in blood are polyethylene terephthalate (PET) and polyurethane (PU).(1, 40) Ming *et al.* detected PET, PA-66 (Nylon-66), PVC, and PE in arterial samples, with PET constituting the highest proportion at 73.70 %. This finding aligns with previous reports identifying PET as the most prevalent MP component in blood.(34, 40) PET, as an emerging plastic raw material in recent years, is widely used in the food, clothing, and packaging industries.(41, 42) In earlier studies focusing on aortic MPs, research by Dong et al. showed that PS-MPs could induce mild vascular calcification in rats.(43) Wang et al. indicated that co-exposure to MPs and lead could induce excessive ROS production in aortic smooth muscle cells, impairing mitochondrial function. However, the researchers used polystyrene (PS) and lead, rather than polyethylene terephthalate (PET), which is the most abundant MP found in the aorta.(10) Research on the toxicological effects of PET-MPs specifically on the aorta remains lacking.

This study investigates the roles of endoplasmic reticulum stress and endothelial glycocalyx damage in PET-MPs-induced aortic injury. Using a co-culture model of HAECs and HASMCs, the research further elucidates the effects of PET-MPs on both aortic endothelial cells and aortic smooth muscle cells. The findings offer novel insights for further elucidating the mechanisms underlying PET-MPs-induced aortic injury.

## 2. Materials and methods

### 2.1 Chemicals

Polyethylene glycol terephthalate MPs were purchased from SuyuanSujiao (China). Nitric acid was purchased from DaMao Chemical Reagent Co., LTD (China). Pentobarbital sodium was purchased from Sigma-Aldrich (Germany). Other reagents used in our study can be found in (Supplementary table 1).

### 2.2 Human samples

Ethical approval for this research was granted by the Clinical Research Ethics Committee of Shanghai General Hospital (YLK2023387), with explicit written consent obtained from all participants prior to the study. Aortic tissue and peripheral blood samples were sourced from sixteen patients diagnosed with abdominal aortic aneurysms. These patients did not exhibit any known genetic disorders. (Supplementary figure 1 and Supplementary table 2).

### 2.3 Animals and treatment

Eight-week-old male Sprague Dawley (SD) rats were sourced from the Experimental Animal Center of Shanghai General Hospital. All animal procedures received approval from the hospital’s Clinical Center Laboratory Animal Welfare & Ethics Committee (2024AW074) and adhered strictly to the National Research Council’s guidelines for laboratory animal care and use. Using drinking water as the method for administering PET-MPs. 21 rats were randomly divided into 4 groups : Control group (deionized water, n=6), low-concentration group (1.0 mg/L PET-MPs suspension, n=5), medium-concentration group (10 mg/L PET-MPs suspension, n=5), and high-concentration group (100 mg/L PET-MPs suspension, n=5). The dosages of PET-MPs were determined based on effective concentrations mentioned in previous studies. Following an 8-week exposure period, all the rats in the above groups were anesthetized with 2 % pentobarbital sodium, and aorta were collected. A range of pathological examinations were conducted to investigate changes in the aortic structure.

### 2.4 Cell culture and treatment

The experiment was performed using immortalized human aortic smooth muscle cells (HASMCs) and immortalized human aortic endothelial cell (HAECs), both obtained from Shanghai ZhongQiaoXinZhou Biotechnology Co., Ltd. (Shanghai, China). Cells were cultured in HASMC complete medium and HAECs complete medium (ZhongQiaoXinZhou Biotechnology Co., Ltd, Shanghai, China), supplemented with 10 % fetal bovine serum (ZhongQiaoXinZhou Biotechnology Co., Ltd, Shanghai, China) and 1 % penicillin/streptomycin (ZhongQiaoXinZhou Biotechnology Co., Ltd, Shanghai, China), and maintained at 37 °C with 5 % CO_2_. HAECs were exposed to PET-MPs at concentrations of 200 μg/ml and 500 μg/ml to investigate the effects of PET-MPs. To determine whether HAECs contribute to the phenotypic transformation of HASMCs, we constructed a conditioned medium cell model. PET-MPs stimulated HAECs with 200 and 500 μg/ml, respectively. Subsequently, the supernatant of the culture medium was collected and used to stimulate HASMCs.

### 2.5 Detection of MPs in blood samples and tissues of patients

Following the method described by Zhang et al. (34), aortic tissue samples were collected in glass containers. Three volumes of nitric acid (68 %) were added to the samples to dissolve the aortic tissues. The samples were then placed on a shaker and agitated at 120 rpm for three days. After complete dissolution of aortic tissues, filtered ultrapure Milli-Q® water was added to the sample vials to adjust the solution pH to 1-2 for subsequent analysis. After complete evaporation of the liquid, the composition of MPs was further analyzed using pyrolysis gas chromatography-mass spectrometry (Agilent Technologies, USA)

### 2.6 Ultrastructure observation

After anesthetizing the rats, aortic tissues were immediately excised and immersed in electron microscope fixative solution. The tissues were then dissected into 1 mm^3^ fragments. Following a 2-hour fixation period at room temperature, the samples were transferred to 4 °C for storage. Used the Leica UC7 ultrathin microtome (Leica, Germany) to cut them into sections of 70-90 nm. Finally, all the slices were subjected to negative staining and observed under a transmission electron microscope (HITACHI, Japan).

### 2.7 Elastic van Gieson (EVG) staining

Aortic tissues underwent fixation, dehydration, and paraffin embedding to generate tissue blocks. Sections of 4 μm thickness were then prepared. For visualization of elastic fibers via EVG staining, sections were immersed in the EVG working solution and incubated at 37 °C for 30 minutes. Subsequent differentiation employed a 2 % ferric chloride solution (10 seconds), followed by counterstaining with Van Gieson’s solution (15 seconds). Sections were then rinsed in absolute ethanol, mounted using neutral resin, and finally subjected to microscopic examination and qualification.

### 2.8 Immunofluorescence

Both aortic tissue sections and treated cells underwent fixation in 4 % paraformaldehyde and subsequent paraffin embedding. Following dewaxing and rehydration, antigen retrieval was performed. Next, the paraffin-embedded aortic sections or treated cells were permeabilized using 0.2 % Triton X-100. Endogenous peroxidase activity was then blocked by incubation in 3 % bovine serum albumin, before supplementation with Wheat Germ Agglutinin (1:100; Thermo, USA), Syndecan 1 Recombinant Rabbit Monoclonal Antibody (1:100; HUABIO, China),Beta Actin Monoclonal antibody (1:100; Proteintech, USA), Calponin Recombinant Rabbit Monoclonal Antibody (1:100; HUABIO, China), alpha smooth muscle Actin Recombinant Rabbit Monoclonal Antibody (1:100; HUABIO, China), Recombinant Human CD31 Protein (1:100; Servicebio, China). Samples were incubated with primary antibody overnight at 4 °C. Following this, the corresponding secondary antibody was applied and incubated for 1 hour at ambient temperature. Subsequent fluorescent labeling utilized Cy3, Cy5, and FITC dyes (Servicebio, Wuhan, China). Cell nuclei were counterstained with DAPI (Servicebio, Wuhan, China). Finally, the sections or cell slides were imaged using a confocal microscope(TCS-SP8, German), and the resulting fluorescent signals were analyzed and quantified.

### 2.9 Western Blot

Cells were collected and immersed in RIPA lysis buffer containing protease inhibitors. An ultrasonic homogenizer was utilized to grind the samples. The samples were then centrifuged at 12,000 rpm for 20 minutes at 4 °C. The supernatant was transferred into an EP tube, added loading buffer, and incubated at 100 °C for 10 minutes. BCA assay was used to quantify the protein concentration. Proteins were separated by SDS PAGE and subsequently transferred onto a PVDF membrane. Primary antibodies were incubated overnight target Syndecan 1 Recombinant Rabbit Monoclonal Antibody (1:1000; HUABIO, China), ICAM-1/CD54 Rabbit Polyclonal Antibody (1:1000; Proteintech, USA), CD54/ICAM-1 Antibody (1:1000;Cell Signaling, USA), NF-κB p65 Rabbit mAb (1:1000; selleck, USA), Phospho-NF-κB p65 (Ser536) Rabbit mAb (1:1000; selleck, USA), XBP1 Recombinant Rabbit Monoclonal Antibody (1:1000; HUABIO, China), ATF4 Recombinant Rabbit Monoclonal Antibody (1:1000; HUABIO, China), Calponin Recombinant Rabbit Monoclonal Antibody (1:1000; HUABIO, China), alpha smooth muscle Actin Recombinant Rabbit Monoclonal Antibody (1:1000; HUABIO, China),GAPDH Rabbit pAb (1:2000; ABclonal, USA),β-actin (1:2000; Proteintech, USA). Following a 1-hour incubation with species-matched secondary antibodies, target protein bands were visualized via chemiluminescent detection. Band intensity quantification was subsequently performed using Image J software.

### 2.10 Quantitative Polymerase Chain Reaction (qPCR) Analysis

qPCR analysis: RNA extraction from HAECs and HASMCs. mRNA was reverse transcribed into cDNA using All-in-one RT SuperMix (Vazyme, China). qPCR was performed using ChamQ Universal SYBR qPCR Master Mix (Vazyme, China) and with the aid of Applied Biosystems ViiA7 real-time fluorescence quantitative PCR instrument (Thermo Fisher, USA). The standardized genes to be tested were calculated using the 2^−ΔΔCT^ method. The mRNA primer sequences used in this study (synthesized by Qingke Biosynthesis) can be found in supplementary information (Supplementary table 3).

### 2.11 Cell viability assay

The CCK-8 assay kit (Servicebio, Wuhan, China) was used to measure the viability of HAECs and HASMCs. The cells were seeded in a 96-well plate at a density of roughly 5,000 cells per well in 100 μL of culture medium and incubated at 37 °C with 5 % CO_2_ for 24 hours. Following treatments, the cells were further incubated at 37 °C with 5 % CO_2_ for 24 hours. After adding 10 μL of CCK8 solution was added to each well, the plate was incubated at 37 °C with 5 % CO_2_ for 1 hour until there was a discernible color shift. The OD value at 450 nm was measured using a microplate reader.

### 2.12 ROS detection

To determine the production of ROS, the Reactive Oxygen Species Assay Kit was used (Servicebio, Wuhan, China). Cells in good condition were selected for detection. The original culture medium was removed, and the cells were washed twice with PBS (Phosphate Buffered Saline). DCFH-DA working solution was added to cover adherent cells in the culture dish, followed by incubation in a 37 °C, 5 % CO₂ incubator protected from light for 45 minutes. After incubation, the working solution was aspirated, and cells were washed twice with PBS. Finally, the cells were covered with PBS. Labeled cells were directly observed under an inverted fluorescence microscope using 488 nm excitation and 525 nm emission wavelengths. Acquired images were subsequently imported into ImageJ software for fluorescence intensity quantification.

### 2.13 Bulk RNA-seq analysis

Rat aortic RNA and HAECs RNA was isolated using Trizol reagent (Invitrogen, CA, USA). Tissue homogenization employed a Servicebio high-speed homogenizer at 4 °C (60 Hz, 3 minutes). RNA integrity was assessed via the RNA Integrity Number on an Agilent 2100 Bioanalyzer (Agilent Technologies, Santa Clara, CA, US). Further purification and quality control utilized an RNAClean XP Kit (Beckman Coulter, CA, USA) combined with an RNase-Free DNase Set (QIAGEN, GmBH, Germany). The difference of gene expression between groups was analyzed using DESeq2 (1.48.1) package in R. Gene set enrichment analysis (GSEA) were performed using fgsea R package and single sample GSEA score of endothelial glycocalyx were calculated based on our in-house gene sets (Supplementary table 4) using GSVA package in R. The transcriptome sequencing results of HAECs and Rat Aorta were found in Supplementary table 5 and Supplementary table 6.

### 2.14 Olink secretome analysis

Following cell culture, the supernatant was harvested and aliquoted into 1.5 ml centrifuge tubes. A refrigerated centrifuge (maintained at 2–8 °C) was employed to pellet cellular debris by centrifugation at 1000–2000 × g for 10 minutes. The resulting supernatant was carefully transferred to new 1.5 ml tubes after discarding the pellet, and subsequently stored at −80 °C. For downstream analysis, 1 μL of this preserved supernatant served as the input material.

Protein quantification was performed using the Proximity Extension Assay (PEA) on the Olink platform (https://www.olink.com). This highly multiplexed immunoassay utilizes validated antibody pairs, each conjugated to unique DNA oligonucleotides, which specifically bind to their target proteins in biological samples. The core principle involves the binding of matched antibody-DNA conjugates to their respective protein targets. When these antibodies bind in close proximity, their attached oligonucleotides hybridize and are extended by DNA polymerase, generating a unique DNA barcode reporter molecule. This barcode is then amplified via PCR and quantified using Next-Generation Sequencing (NGS).

Specifically, the assay involved incubating samples with 372 distinct antibody-DNA pairs for 16-24 hours to facilitate target binding and oligonucleotide hybridization. The subsequent extension, PCR amplification, and NGS quantification steps were carried out. NGS was conducted on an Illumina NovaSeq 6000 platform, and the generated data (Normalized Protein eXpression, NPX) were processed using the Olink NPX Explore Software.

The experimental procedures encompassing incubation, oligonucleotide extension/ligation, and detection for the Olink PEA were outsourced to and executed by Sinotech Genomics Co., Ltd. (Shanghai, China). Differential analysis of relative protein expression levels between experimental groups was conducted utilizing the limma package within the R statistical environment.

### 2.15 Statistical analyses

Continuous variables are expressed as mean ± SEM from a minimum of three independent replicates. Normality distribution was verified via Shapiro-Wilk testing. Two-group comparisons employed unpaired Student’s t-test for parametric data; alternatively, the Mann-Whitney U-test addressed non-parametric datasets. Multi-group analyses utilized one-way ANOVA with post hoc testing for normally distributed variables, whereas non-normally distributed data underwent Kruskal-Wallis testing with Dunn’s correction. Statistical procedures were executed in GraphPad Prism 8.0, with significance thresholds set at P < 0.05.

## 3. Results

### 3.1 Distribution of microplastics in human aorta and characterization of PET-MPs

We collected aortic and blood samples from patients with aortic aneurysms. The plastic composition within these samples was analyzed using pyrolysis-gas chromatography to verify the relative abundance of 4 kinds of common microplastic in the human aorta. The results indicated a relatively higher proportion of microplastic in the human aorta than others (Fig 1D-F and Supplementary table 7). Additionally, we characterized the properties of the PET-MPs used in the simulated microplastic exposure in rats. Their morphology under scanning electron microscopy is shown in Fig 1B and C, with an average diameter of 6.0402 μm, classifying them as micron-sized particles (Supplementary figure 2A).

**Figure 1.**
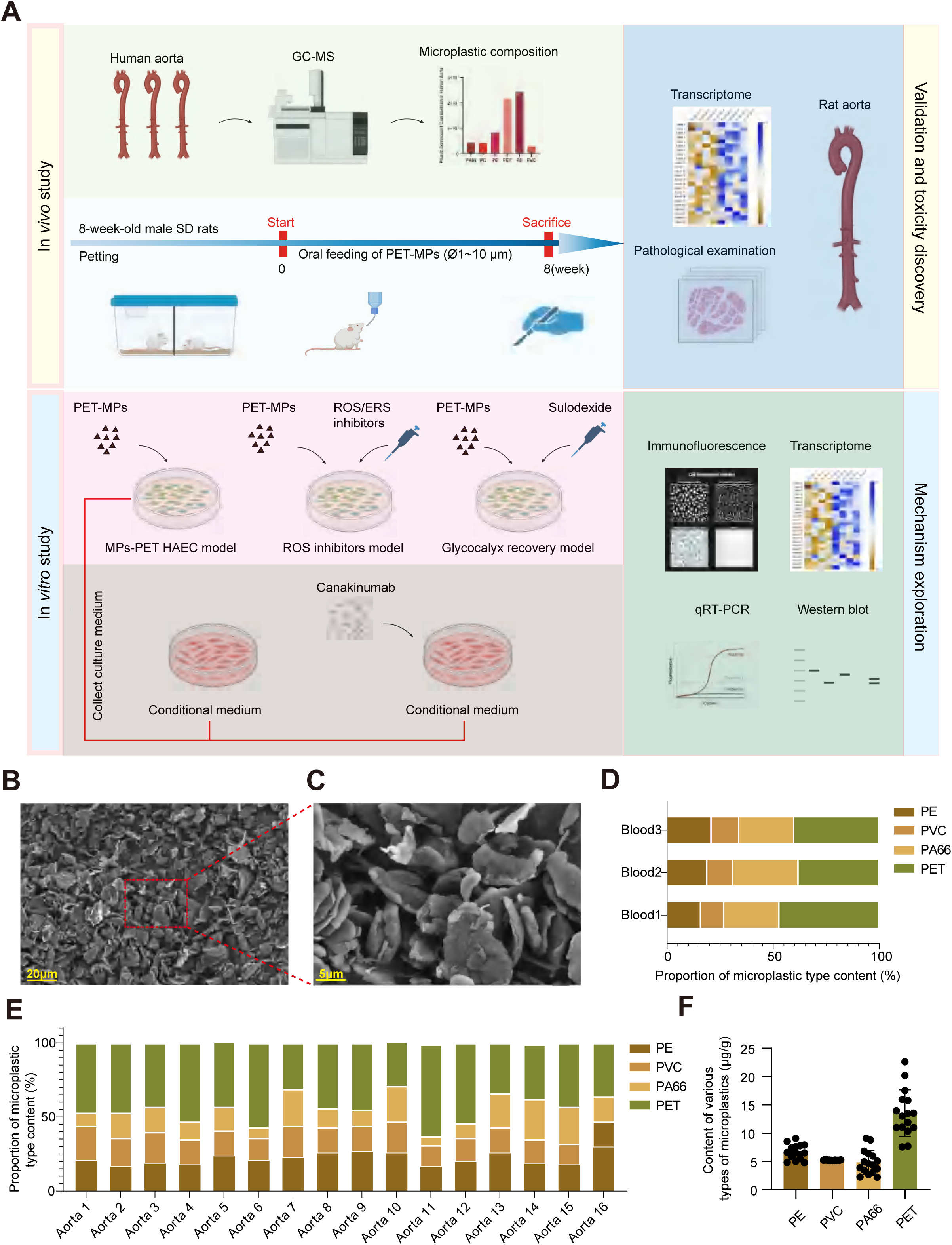

### 3.2 PET-MPs deposition in aortic endothelial cells causes early structural impairment after chronic exposure in rats

The morphology of the rat aorta and the content of PET microplastics are shown in Fig 2A,B. Pathological examination of rat aortas revealed that stimulation with different concentrations of PET-MPs did not induce significant changes in aortic wall thickness (Fig 2A, D). Notably, the morphology of the aortic elastic fibers exhibited notable alterations (Fig 2A, C). Under normal physiological conditions, aortic elastic fibers display a coiled configuration to maintain aortic elasticity. In this study, exposure to varying concentrations of PET-MPs caused the elastic fibers to lose their original morphology to varying degrees (Fig 2A, C). Transmission electron microscopy observations revealed that PET-MPs primarily deposited within the endothelial cells of the rat aorta; no PET-MPs particles were detected within the elastic fiber layer (Fig 2E, F).

**Figure 2.**
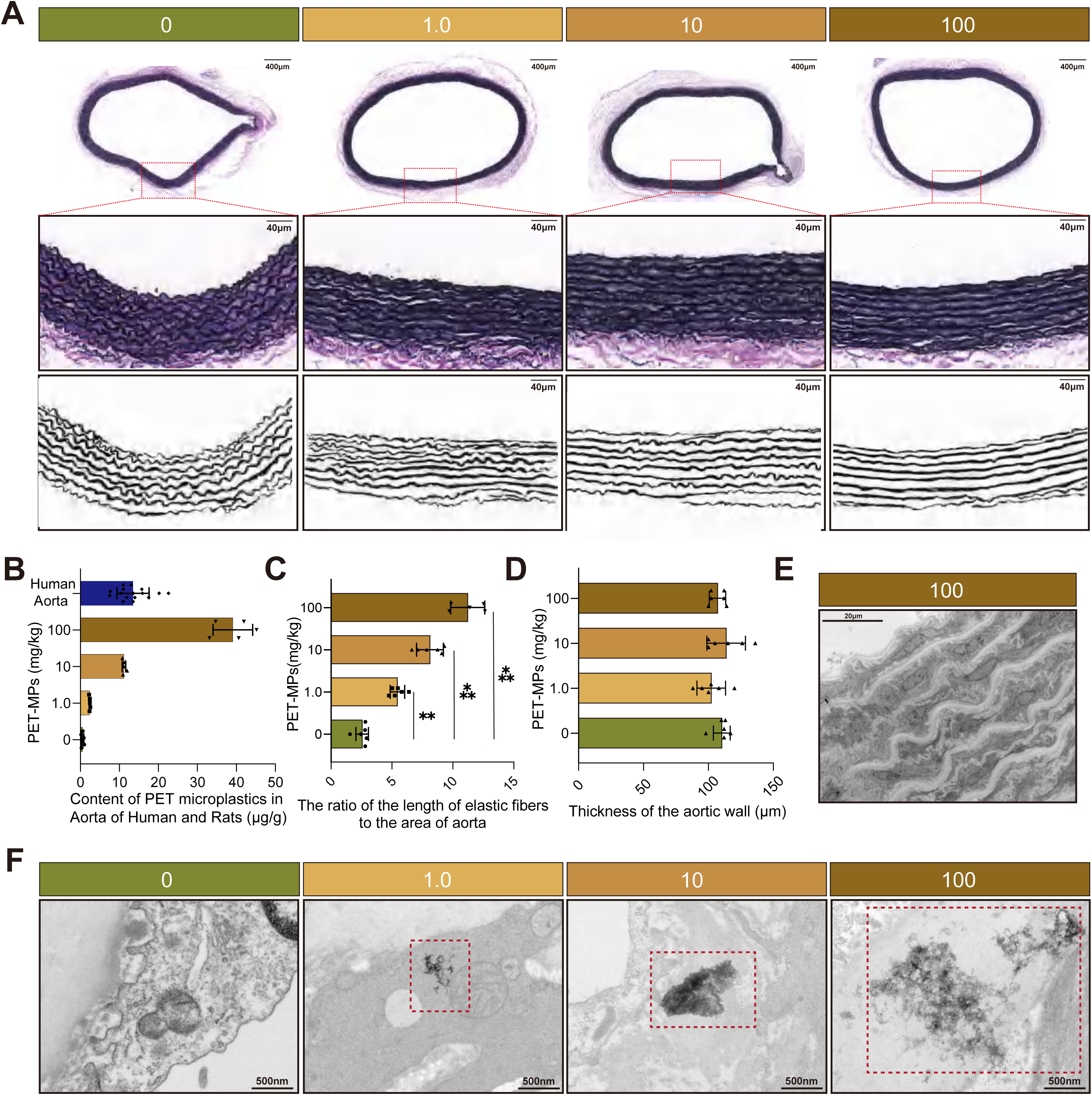

These results suggest that PET-MPs likely first target rat aortic endothelial cells, subsequently affecting rat aortic smooth muscle cells—which constitute the aortic elastic media—potentially via cell-cell interactions.

### 3.3 PET-MPs disrupt the endothelial glycocalyx which contributes to early aortic injury

To investigate the toxicological mechanism underlying PET-MPs-induced aortic alterations in rats, we performed transcriptomic analysis (Fig 3A, B). Gene Set Enrichment Analysis (GSEA) of rat aortas showed alterations in pathways related to extracellular matrix degradation and endothelial glycocalyx homeostasis (Fig 3A, B). Furthermore, reduced phenotypic score of glycocalyx biosynthesis and luminal glycocalyx correlated inversely with increased curvature of the aortic elastic fibers, indicating early structural impairment (Fig 3B).

**Figure 3.**
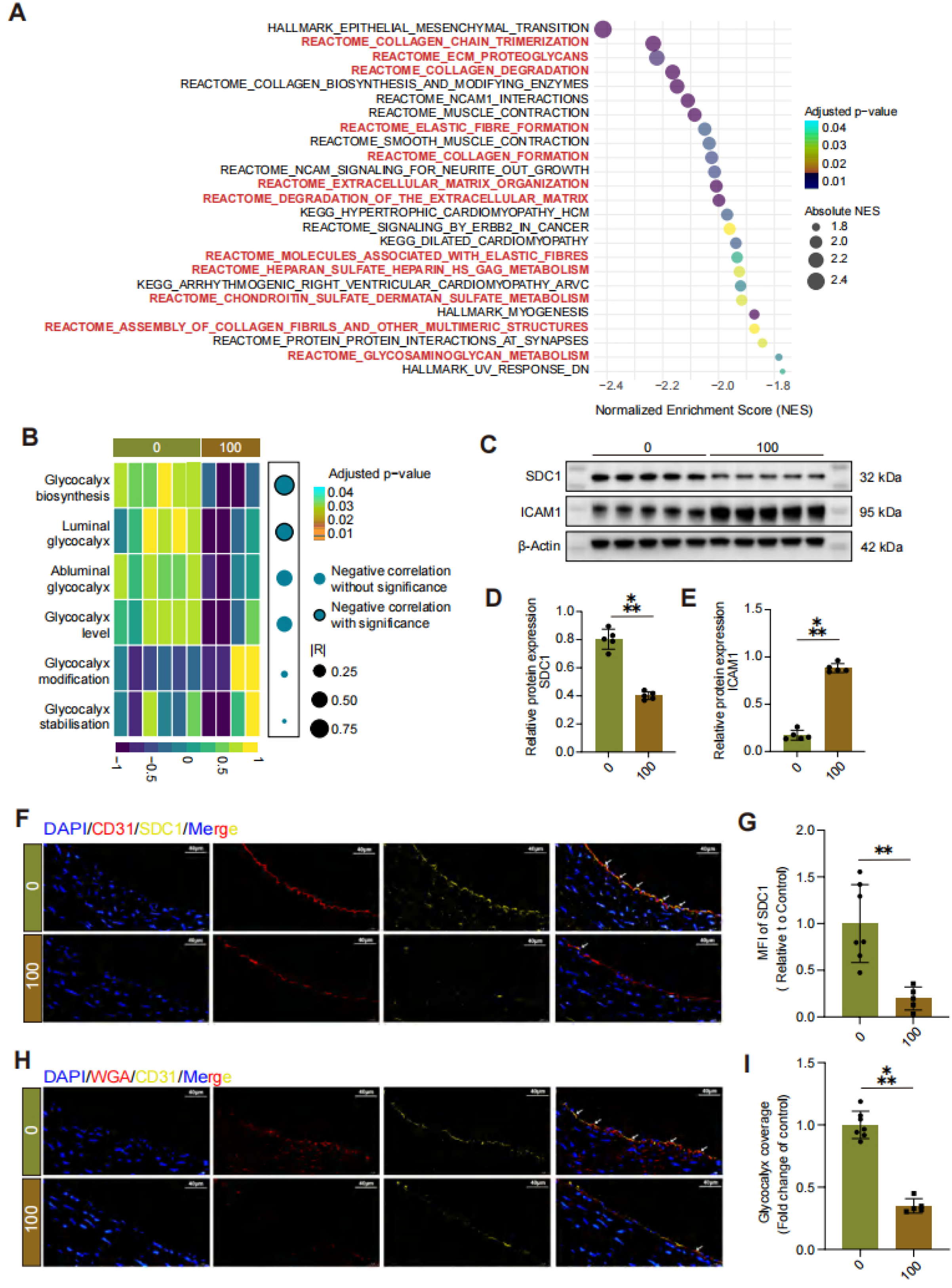

Multiple studies have established the crucial role of the endothelial glycocalyx in maintaining vascular stability.(11–16) Damage to the glycocalyx impairs several key functions of the endothelium. Western blot analysis demonstrated that rats exposed to 100 mg/L PET-MPs exhibited increased ICAM1 protein expression and decreased SDC1 protein expression in the aorta, indicating endothelial glycocalyx damage and endothelial activation (Fig 3C-E).

Wheat Germ Agglutinin (WGA) is a non-immunogenic sugar-binding protein that specifically recognizes mono- or oligosaccharide structures in glycoproteins and glycolipids on the cell surface within the vascular lumen. We utilized WGA to specifically recognize and bind to the glycocalyx structure on the luminal surface of rat aortas for observation and semi-quantitative assessment. To further investigate whether the rat aortic endothelial glycocalyx was damaged, we employed immunofluorescence to visualize key glycocalyx components, SDC1 and WGA. The results showed that compared to control rats, rats exposed to 100 mg/L PET-MPs exhibited decreased fluorescence intensity for both SDC1 protein and WGA in the aortic endothelium (Fig 3F-I).

In summary, these findings indicate that PET-MPs induce aortic endothelial damage via glycocalyx degradation, thereby triggering early pathological remodeling characterized by elastic fiber impairment .

### 3.4 PET-MPs suppress glycocalyx homeostasis in human aortic endothelial cells

Given the pivotal role of the rat aortic endothelium in aortic injury, we established a PET-MPs exposure model using Human Aortic Endothelial Cells (HAECs) to simulate the *in vivo* stimulation process. CCK-8 assays were used to determine appropriate microplastic concentrations for stimulating HAECs: low concentration (MPs+) at 200 μg/ml and high concentration (MPs++) at 500 μg/ml (Supplementary figure 2B,C). After 24 hours of PET-MPs stimulation, culture supernatant was collected, and the remaining cells underwent transcriptome sequencing. Similarly, transcriptome sequencing results revealed that compared to the control group, the endothelial glycocalyx related pathway was significantly downregulated in the PET-MPs exposed HAECs, suggesting damage to the glycocalyx structure (Fig 4A, B). Western blot analysis also confirmed a significant decrease in SDC1 protein levels in PET-MPs exposed HAECs (Fig 4C).

**Figure 4.**
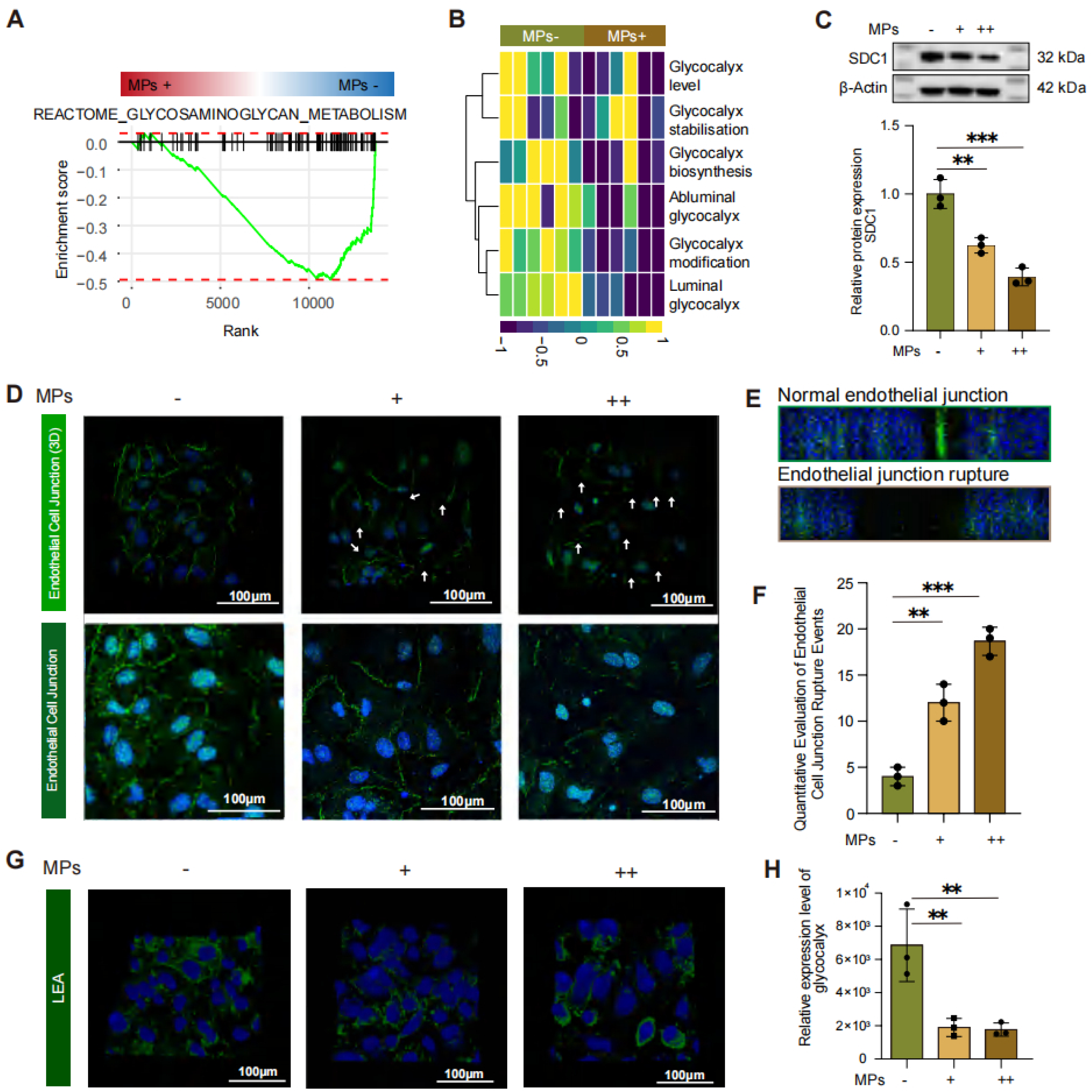

β-catenin is a core protein of endothelial adherent junctions, maintaining vascular barrier integrity by anchoring other proteins to the cytoskeleton. Decreased β-catenin expression or abnormal localization directly leads to increased endothelial permeability. Our immunofluorescence cell staining results demonstrated that stimulation with different concentrations of PET-MPs reduced β-catenin expression in HAECs and caused multiple breaks and losses in β-catenin connections between endothelial cells, indicating potential impairment of the HAECs barrier function (Fig 4D-F).

Lycopersicon Esculentum Lectin (LEA) is a non-immunogenic sugar-binding protein that specifically recognizes mono- or oligosaccharide structures in cell surface glycoproteins and glycolipids. We employed LEA to specifically recognize and bind to the glycocalyx structure on HAECs surfaces for observation and semi-quantitative detection. Immunofluorescence cell staining results confirmed a decrease in glycocalyx content on HAECs stimulated with different concentrations of PET-MPs (Fig 4G-H).

In conclusion, these results indicate that PET-MPs exposure disrupts endothelial barrier function and damages the glycocalyx in HAECs, mirroring the injury mechanisms observed in the rat aortic endothelium.

### 3.5 PET-MPs induce the ER stress-ROS axis which mediates glycocalyx disruption

To further elucidate the pathways through which PET-MPs damage the endothelial glycocalyx in HAECs, we analyzed the transcriptome sequencing results from the PET-MPs exposed cell model. GSEA revealed significant upregulation of the pathway associated with Endoplasmic Reticulum (ER) stress activation (Fig 5A-B). ATF4 and XBP1 are core regulators of ER stress, and changes in their expression levels directly reflect the state of ER oxidative stress. Stimulation with different concentrations of PET-MPs induced a dose-dependent increase in both the transcriptional and translation levels of ATF4 and XBP1 in HAECs (Fig 5C-G).

**Figure 5.**
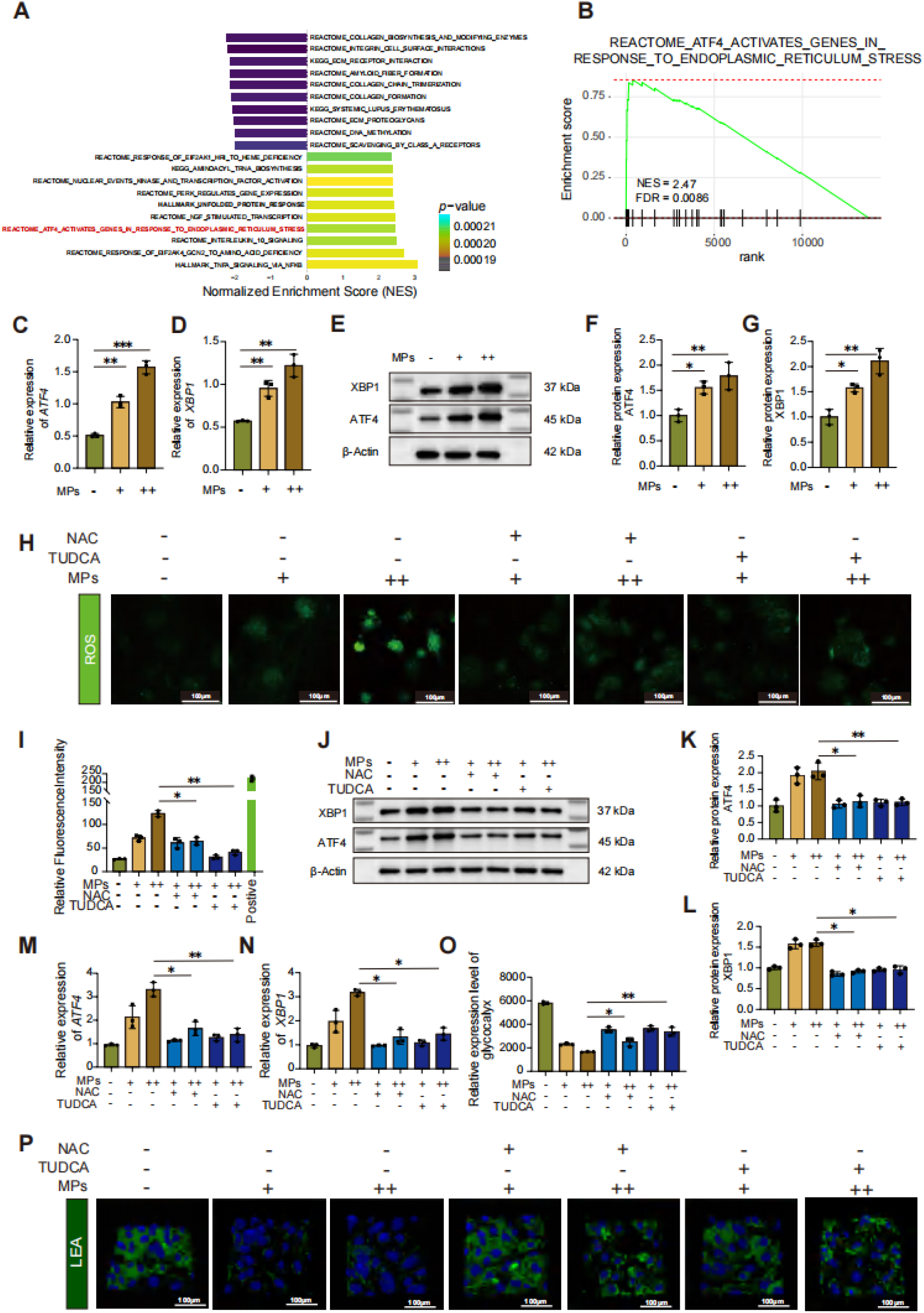

To investigate the relationship between ER oxidative stress and HAECs glycocalyx damage, we treated the cell model with two inhibitors: N-acetylcysteine (NAC) and tauroursodeoxycholic acid (TUDCA). Appropriate treatment concentrations were determined using the CCK-8 assay (Supplementary figure 2D,E). We employed 2’,7’-dichlorodihydrofluorescein diacetate (DCFH-DA) for semi-quantitative fluorescence detection of ROS. The results showed that PET-MPs exposure induced a dose-dependent increase in DCFH-DA fluorescence intensity. Compared to HAECs directly exposed to PET-MPs without NAC or TUDCA pretreatment, HAECs pretreated with NAC or TUDCA exhibited significantly lower DCFH-DA fluorescence intensity (Fig 5H,I). This indicates that NAC and TUDCA can both reduce the level of oxidative stress in HAECs exposed to PET-MPs. Consistent with this trend, ATF4 and XBP1 expression levels also decreased at both transcriptional and protein translation levels following pretreatment (Fig 5J-N). Subsequently, LEA staining was again used for semi-quantitative fluorescence detection of glycocalyx content in the different HAECs groups. The results showed that pretreatment with NAC and TUDCA, which reduced ER oxidative stress levels, also partially alleviated the degree of glycocalyx structural damage (Fig 5O,P).

In summary, this series of experiments demonstrate that PET-MPs induces ER stress in HAECs, leading to increased ROS production, which in turn mediates the disruption of endothelial glycocalyx.

### 3.6 Glycocalyx loss by PET-MPs activates NF-κB-driven endothelial inflammation

The NF-κB pathway is rapidly activated during inflammation, regulating the expression of numerous key genes involved in inflammation, immune responses, and cell survival. In our study, transcriptome sequencing of the PET-MPs exposed cell model revealed that PET-MPs exposure significantly activated and upregulated the NF-κB pathway (Fig 6A). Western blot analysis corroborated this finding, showing upregulated ICAM1 expression and an increased pP65/tP65 ratio in HAECs exposed to PET-MPs, indicating endothelial activation and an inflammatory state (Fig 6B-E). Furthermore, we observed upregulated transcriptional levels of inflammatory cytokines such as IL-1β, IL-6, and IL-8 (Fig 6F-H). These results collectively indicate that PET-MPs exposure induces an inflammatory state in HAECs.

**Figure 6.**
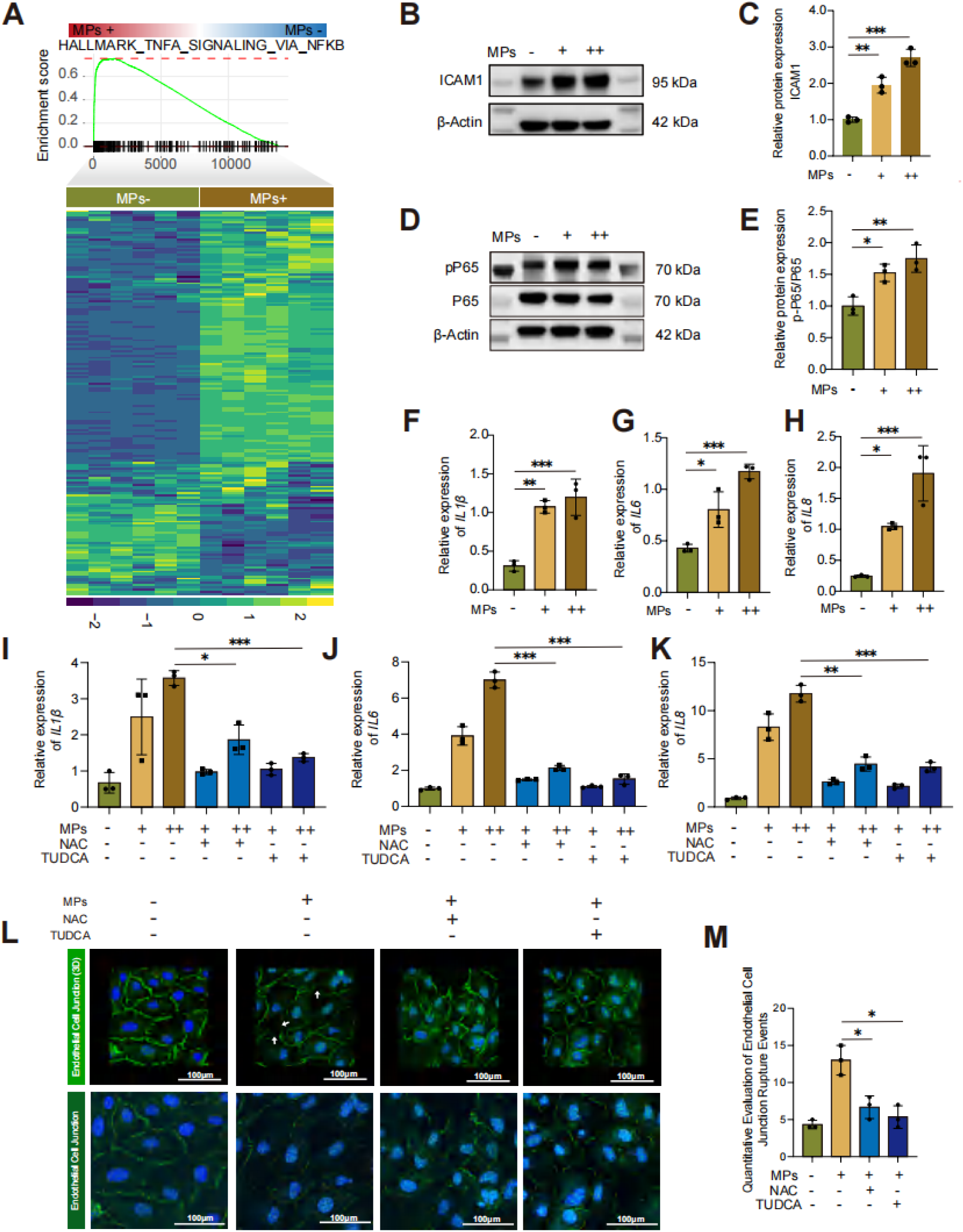

Does the activation of ER oxidative stress, while causing glycocalyx damage, also contribute to the onset of the endothelial inflammatory state? To explore this question, we again pretreated cells with oxidative stress inhibitors NAC and TUDCA. The results showed that inhibiting oxidative stress damage improved the inflammatory state of the cells. Compared to the PET-MPs exposed group without inhibitors, the transcriptional expression levels of inflammatory cytokines IL-1β, IL-6, and IL-8 were downregulated following inhibitor pretreatment (Fig 6I-K). Additionally, the endothelial junction damage caused by PET-MPs exposure was partially restored after inhibiting oxidative stress damage with NAC and TUDCA (Fig 6L-M).

In conclusion, these results suggest that the damage induced by ER oxidative stress not only triggers disruption of the endothelial glycocalyx structure but also promotes an inflammatory state in endothelial cells, leading to upregulated expression of inflammatory cytokines.

### 3.7 PET-MPs trigger secretome alterations in human aortic endothelial cells and phenotypic modulation of human aortic smooth muscle cells

To identify which inflammatory factors might be secreted by HAECs in response to PET-MPs exposure, we collected culture supernatants from the PET-MPs exposed cell model and analyzed the relative levels of inflammatory factors using Olink assay technology (Fig 7A). The Olink assay results showed: 48 proteins were significantly upregulated in the Medium MPs group compared to Control, 52 proteins were significantly upregulated in the High MPs group compared to Control, and 15 proteins were significantly upregulated in High MPs compared to Medium MPs. After intersecting these comparisons, 5 proteins were selected: IL-1β, IL-32, DFFA, EGLN1, and HCLS1 (Fig 7B-E) (Supplementary figure 2G-I). Among these, the upregulation of both IL-1β and IL-32 (which is IL-1β dependent, Fig 7B) indicates a broad inflammatory activation state. However, IL-1β warrants particular attention due to its central role in endothelial cell activation; it directly drives key inflammatory responses and smooth muscle phenotype changes closely related to the pathological processes under investigation.(44, 45) Therefore, this study focused on IL-1β as a key inflammatory cytokine potentially responsible for inducing phenotypic changes in HASMCs.

**Figure 7.**
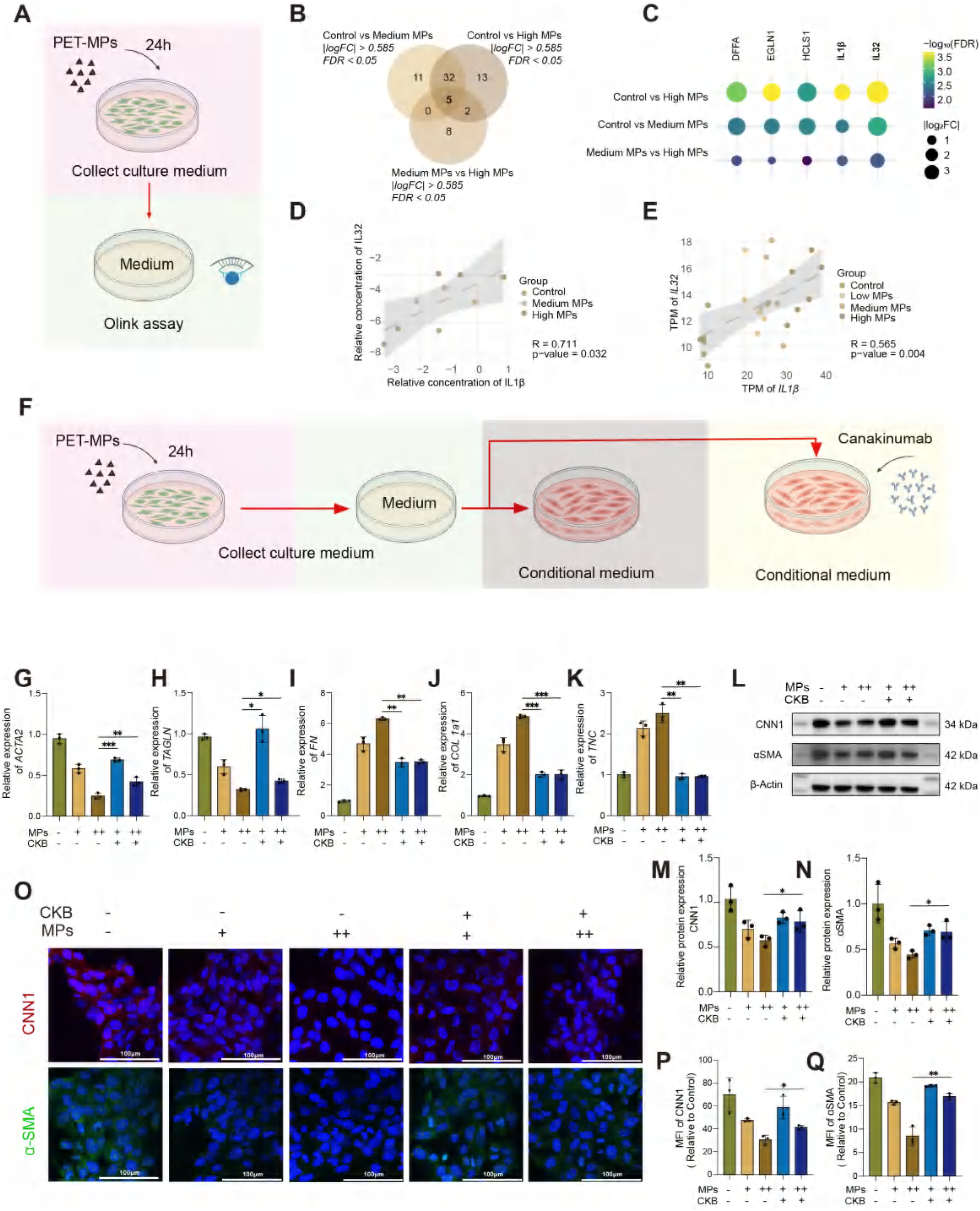

We established a conditioned medium cell model for HASMCs: using the supernatant from the PET-MPs exposed HAECs cell model to stimulate HASMCs. Canakinumab (an IL-1β monoclonal antibody) was used to attempt blocking this stimulatory effect (Fig 7F). The results showed that at both transcriptional and protein translation levels, HASMCs stimulated with the conditioned medium underwent a phenotypic switch towards a synthetic phenotype. Intervention with Canakinumab attenuated the degree of this switch towards a synthetic phenotype in HASMCs (Fig 7G-N). Immunofluorescence results similarly confirmed this; stimulation with conditioned medium reduced the expression of contractile phenotype markers in HASMCs. However, this decrease was partially mitigated following Canakinumab intervention (Fig 7O-Q).

We suggested that IL-1β, secreted by HAECs in an inflammatory state, is a key factor contributing to the phenotypic modulation of HASMCs, potentially initiating the structural degenerative changes observed in the aortic elastic fibers.

### 3.8 Sulodexide restores glycocalyx and ameliorates endothelial cell activation induced by PET-MPs

Given the significant upregulation of ER oxidative stress observed in this study and its damaging effect on the endothelial glycocalyx structure, exploring drugs with glycocalyx-repairing capabilities, such as Sulodexide, may offer new avenues for intervention strategies. Therefore, we administered Sulodexide in the PET-MPs exposed HAECs cell model. The results demonstrated that intervention with Sulodexide partially restored the glycocalyx structure in HAECs exposed to different concentrations of PET-MPs (Fig 8A, B). Western blot analysis also confirmed this restoration (Fig 8C, D).

**Figure 8.**
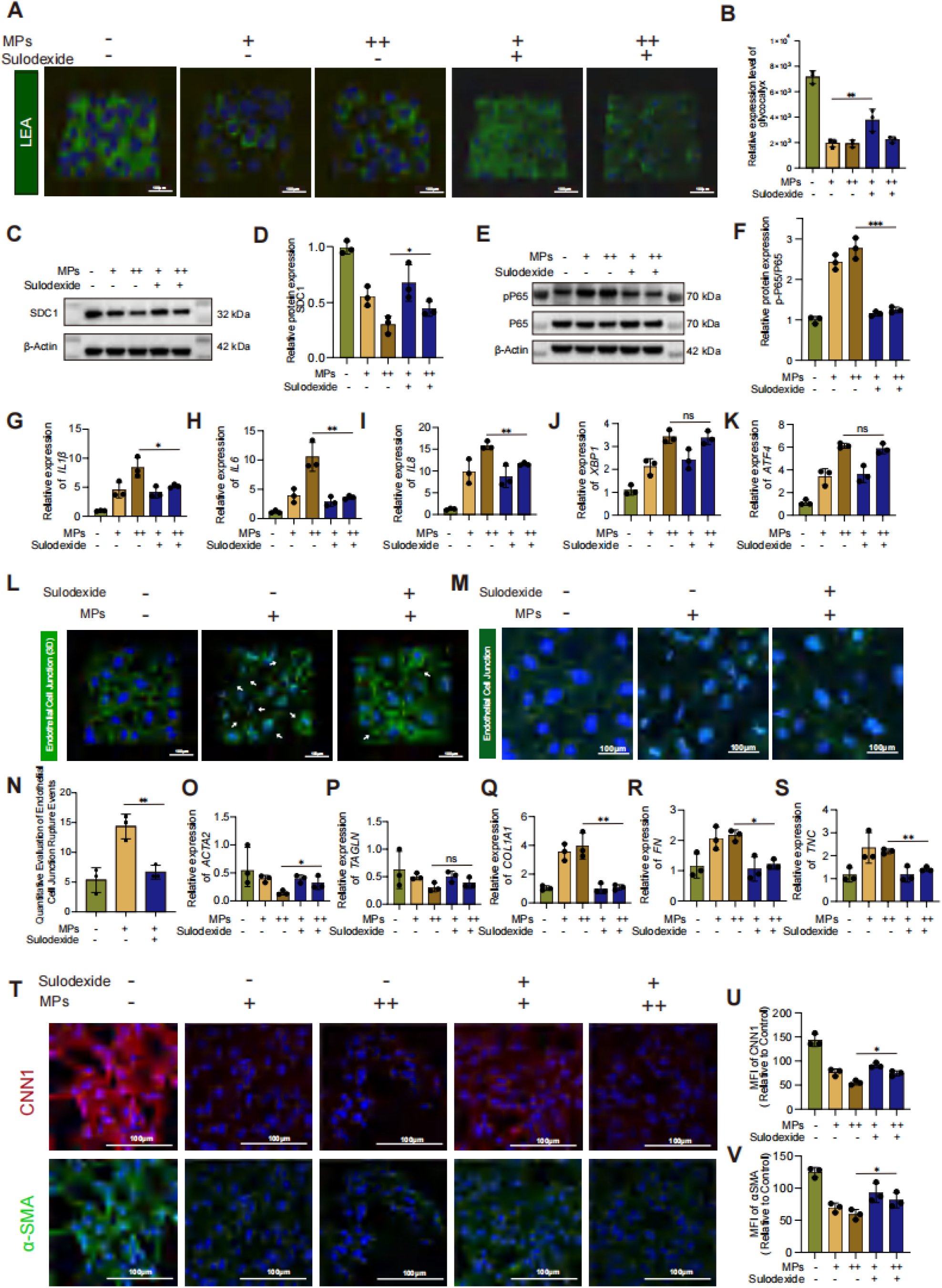

As the glycocalyx structure is a vital component of endothelial cell functional integrity, does its restoration influence the inflammatory state of HAECs? Western blot and qRT-PCR results showed that Sulodexide intervention reduced the PET-MPs-induced increase in the pP65/tP65 ratio (Fig 8E-F). Studies by Yuan et al. also support this finding.(46) Similarly, the expression of inflammatory cytokines IL-1β, IL-6, and IL-8 and ER-Stress-related genes were downregulated (Fig 8G-K). Furthermore, the PET-MPs-induced disruption of endothelial junctions was partially restored following Sulodexide intervention (Fig 8L-N).

We then reused the HASMC conditioned medium cell model, pretreating HAECs with Sulodexide before collecting the supernatant for HASMC stimulation. The results showed that intervention with Sulodexide on HAECs inhibited the process of phenotypic modulation in HASMCs (Fig 8O-V).

In summary, the glycocalyx, as a critical functional structure of the vascular endothelium, is of paramount importance. Restoration of the endothelial glycocalyx structure can partially improve endothelial function and suppress the endothelial inflammatory state.

### 3.9 PET-MPs exposure signature links to human arterial pathologies

Given that PET-MPs induced aortic remodeling in this study resembles the pathological process of aortic diseases, we defined a ’PET-MPs exposure signature’ (MPS) gene set based on transcriptomic data from PET-MPs exposed endothelial cells, comprising 150 significantly upregulated genes and 150 significantly downregulated genes. (Supplementary table 8) To investigate the potential significance of MPS in arterial diseases, we applied it to gene set enrichment analysis across multiple independent datasets. The results revealed that MP UP signature was significantly enriched in disease phenotype of abdominal aortic aneurysm (Fig 9A-C), thoracic aortic aneurysm (Fig 9D), and aortic dissection (Fig 9E,F) samples. In terms of prognostic association, survival analysis using data from patients who underwent carotid endarterectomy indicated that higher MP UP signature score was significantly correlated with the higher risk of postoperative ischemic events (Fig 9G). Additionally, MP UP signature were enriched in oxidized phospholipids treated aortic endothelial cells and TNF-α treated human umbilical vein endothelial cells (Fig 9H,I). This signature bridges experimental PET-MPs exposure to human disease mechanisms and clinical outcomes.

**Figure 9.**
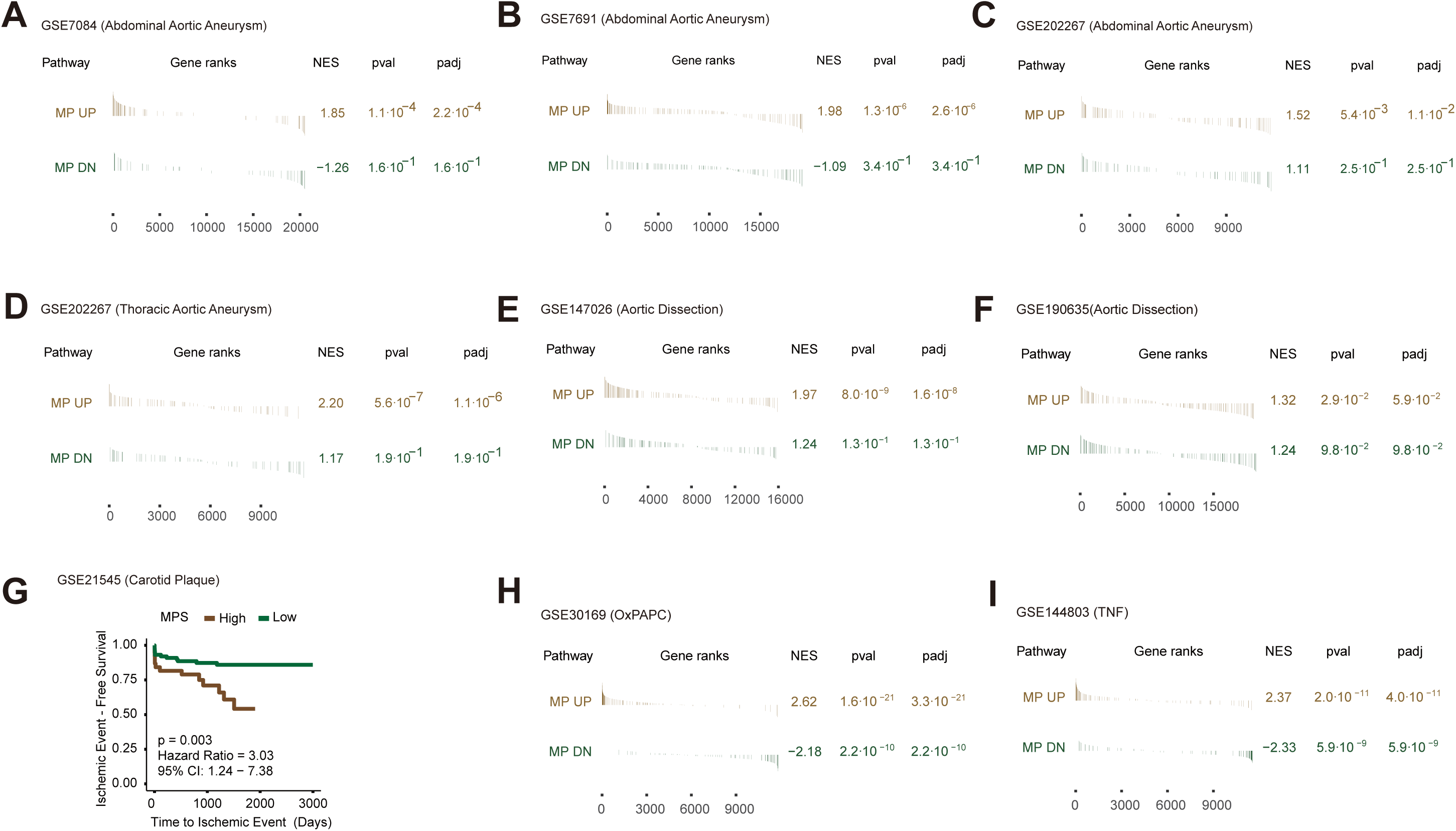

## 4. Discussion

Our current study confirms that the loss of the endothelial glycocalyx plays a pivotal role in the response to polyethylene terephthalate microplastics (PET-MPs). By establishing a chronic PET-MPs exposure model in rats, we found that early structural damage to the aorta is closely associated with disruption of the endothelial glycocalyx. At the cellular level, our data reveal endoplasmic reticulum stress (ERS) as the initiating factor, activating the generation of reactive oxygen species (ROS) and establishing a self-amplifying ERS-ROS loop that mediates glycocalyx degradation. This cascade subsequently triggers endothelial inflammation, compromises endothelial integrity, and alters the endothelial secretome profile, ultimately promoting phenotypic switching of aortic smooth muscle cells. Our study highlights the intimate connection between aortic endothelial cells and smooth muscle cells. Intervention with Sulodexide not only restored the endothelial glycocalyx but subsequently reduced the activated state of the endothelium.

Although the critical role of the endothelial glycocalyx in vascular homeostasis is well recognized (11, 19, 20), our study is the first to identify PET-MPs as a novel environmental trigger causing endothelial glycocalyx injury. The glycocalyx structure was disrupted in the aortic endothelium of rats exposed to PET-MPs. In the human aortic endothelial cell (HAECs) model, we found that PET-MPs stimulation similarly compromised the integrity of the endothelial glycocalyx structure. Notably, glycocalyx loss occurred prior to overt inflammation or smooth muscle phenotypic changes, suggesting its role as a "first responder" to PET-MPs exposure. Furthermore, the degree of downregulation in SDC1 expression, a sentinel for vascular stress, reflects the extent of vascular damage induced by PET-MPs exposure .(47–49)

Pathways leading to glycocalyx damage include oxidative stress, inflammation, proteolytic enzyme activity, and dysregulation of mechanical stress.(50–52) How then do PET-MPs disrupt the aortic endothelial glycocalyx? Using transmission electron microscopy, we observed PET-MPs internalized within rat aortic endothelial cells, which may directly initiate endothelial cell injury. Secondly, transcriptomic analysis of rat aortas revealed a significant upregulation of genes related to ER oxidative stress in rats exposed to high concentrations of PET-MPs compared to controls, suggesting this as a key mechanism of PET-MPs-induced endothelial injury. The endoplasmic reticulum (ER) is a vital membranous organelle in eukaryotic cells responsible for protein synthesis, folding, and secretion. When the ER is subjected to endogenous or exogenous stimuli, its protein folding function becomes disrupted, triggering a cascade of subsequent reactions termed ER oxidative stress.(25, 26) Kaufman *et al.* indicates that ER stress leads to elevated levels of ROS and calcium, causing further damage.(27, 28) Wu *et al.* also demonstrate that ER stress is a major source of ROS.(53)

ROS not only poses a direct threat to glycocalyx but also enhances its proteolysis by activating matrix metalloproteinases (MMPs) and inactivating endogenous protease inhibitors.(29, 30) Furthermore, existing literature reports that ER stress accompanies vascular calcification in mice and rats.(54, 55) NAC is a broad-spectrum antioxidant that directly scavenges ROS and enhances glutathione synthesis, while TUDCA inhibits the source of ER stress by blocking PERK/IRE1 pathway activation, thereby indirectly alleviating ER oxidative stress. Following treatment with TUDCA or NAC, PET-MPs-induced endothelial glycocalyx damage was significantly mitigated. This indicates that, distinct from factors like inflammation or mechanical stress dysregulation, PET-MPs elevate ROS by inducing ER oxidative stress, subsequently damaging the endothelial glycocalyx via oxidative stress. Notably, the inhibitory effect of TUDCA surpassed that of NAC, suggesting that ER oxidative stress may be the primary trigger for endothelial glycocalyx disruption.

Wang *et al.* showed that damage to the endothelial glycocalyx activates the NF-κB pathway, promoting the release of pro-inflammatory cytokines such as IL-6 and TNF-α, which further activate MMPs and heparanase, establishing a vicious cycle of "glycocalyx damage-inflammation-enzymatic degradation".(56–58) Our study elucidates how PET-MPs exploit this damaging pathway to drive degenerative changes in the aortic vascular wall. Transcriptomic analysis of rat aortas revealed significant activation of the NF-κB pathway and upregulated expression of IL-1β and IL-6 in PET-MPs exposed rats, mirroring the phenotype of glycocalyx loss observed under pathological conditions.(46) Additionally, Olink assay results demonstrated that conditioned medium from PET-MPs-exposed endothelial cells was enriched in IL-1β. IL-1β has been shown to induce downregulation of contractile phenotype marker proteins in smooth muscle cells, promoting SMC phenotypic switching—a hallmark of early vascular remodeling.(59, 60) Neutralization of IL-1β effectively prevented this phenotypic modulation, indicating that endothelial-derived inflammatory cytokines play a critical role in initiating downstream changes in the vascular wall, particularly the phenotypic alteration of vascular smooth muscle cells.

Sulodexide is a natural glycosaminoglycan mixture extracted from porcine intestinal mucosa, primarily composed of fast-moving heparin fraction and dermatan sulfate. This drug has garnered attention for its unique endothelial protective effects, particularly its ability to repair damaged endothelial glycocalyx.(61) Furthermore, it protects the vascular endothelium by modulating endothelial permeability and restoring the endothelial glycocalyx.(62) Our results demonstrate that sulodexide directly restores the PET-MPs-impaired glycocalyx in HAECs and can protect endothelial cell function while suppressing inflammatory activation. Sulodexide has previously been used for various vascular-related diseases(63–65), and our findings further highlight its therapeutic potential against PET-MPs-induced vascular injury.

In this study, aortic remodeling induced by PET-MPs primarily involved degenerative changes in the medial elastic fibers, specifically the loss of their characteristic undulations. This feature resembles pathological processes observed in numerous aortic diseases(66, 67). This striking similarity raises a critical question: does PET-MPs exposure recapitulate molecular pathways involved in clinical aortic pathologies? To address this, we developed a "PET-MPs Exposure Signature" (MPS) based on transcriptomic changes in aortic endothelial cells exposed to PET-MPs. When we compared this signature with human aortic disease datasets, GSEA analysis demonstrated significant enrichment of MP-UP-associated genes in patient-derived samples and *in vitro* disease models, reinforcing the pathophysiological relevance of PET-MPs-induced vascular injury. These findings suggest that microplastic exposure may contribute to aortic disease progression by activating conserved pathological pathways, providing a novel environmental link to vascular degeneration. Future research, encompassing further experimental studies and epidemiological investigations, is required to validate this association and elucidate the mechanisms by which PET-MPs exposure participates in the pathological processes of arterial diseases.

Several limitations of current study should be considered. Firstly, the *in vivo* doses (up to 100 mg/L) and in *vitro* concentrations (up to 500 μg/ml) significantly exceed estimated human environmental exposure levels (mg/kg/day). While high doses are often necessary for mechanistic exploration in controlled models, this limits direct extrapolation to human risk assessment; future studies employing environmentally relevant doses and chronic exposure paradigms are crucial. Secondly, we used pristine PET-MPs, whereas aged/weathered MPs in the environment often exhibit altered surface properties and adsorbed pollutants, potentially modifying toxicity. Thirdly, the lack of direct *in vivo* evidence for ERS activation, ROS elevation, and SMC phenotype marker changes weakens the direct translation of the detailed *in vitro* mechanism to the observed pathology. Fourthly, static cell culture neglects physiologically relevant hemodynamic shear stress, a key regulator of endothelial function and glycocalyx stability. Fifthly, potential species differences between rats and humans necessitate caution. Sixthly, assessing only an 8-week endpoint precludes insights into the progression or recovery of lesions. Finally, functional consequences (e.g., aortic stiffness, blood pressure changes) were not measured. Addressing these limitations will be essential to fully understand the pathophysiological relevance and translational potential of our findings.

## 5. Conclusion

In this study, we unveil the endothelial glycocalyx as a critical and sensitive nexus mediating vascular responses to microplastic exposure. By delineating a cascading mechanism where microplastics trigger ER stress and oxidative imbalance leading to glycocalyx erosion, vascular inflammation, and early smooth muscle dysregulation, we provide new mechanistic insights into the vascular consequences of environmental pollutants. Remarkably, the demonstration that glycocalyx integrity can be restored pharmacologically highlights a promising, actionable target for vascular protection. Our findings bridge environmental toxicology and cardiovascular pathophysiology with urgent clinical implications.

## Supporting information

Supplementary table 1: Detailed information on the antibodies, kits and other consumables used in this study.

Supplementary table 2:The basic information of 16 patients.

Supplementary table 3: Primer sequences used in this study.

Supplementary table4: Calculate the gene set required for the endocalyx.

Supplementary table 5: HAECs transcriptome sequencing results.

Supplementary table 6: Rat Aorta transcriptome sequencing results.

Supplementary table 7:Py-GC/MS calibration curves of polymers.

Supplementary table 8:PET-MPs exposure signature.

Supplementary Figure l. Gross morphology of arterial samples

## CRediT authorship contribution statement

Weixue Huo: Writing–original draft, Investigation, Conceptualization. Lushun Yuan: Writing review & editing, Funding acquisition, Conceptualization. Rui Feng: Supervision, Funding acquisition. Sen Wang: Visualization, Methodology, Investigation. Deping Kong: Investigation, Formal analysis, Conceptualization. Jin Qu: Resources, Data curation. Mengwei He: Visualization, Methodology. Zhaoxiang Zeng: Methodology, Investigation, Data curation.

## Ethical approval

Ethical approval for this research was granted by the Clinical Research Ethics Committee of Shanghai General Hospital (YLK2023387) and all animal procedures received approval from the Shanghai General Hospital’s Clinical Center Laboratory Animal Welfare & Ethics Committee (2024AW074)

## Appendices. Supplementary material

Supplementary data associated with this article can be found in Supplementary table 1-8 and Supplementary figure 1-2.

## Data availability

Data will be made available on request.

The original data of transcriptome sequencing in this study: PRJNA1287132, PRJNA1287283

## Abbreviations

AAA: Abdominal Aortic Aneurysm
AD: Aortic Dissection
ATF4: Activating Transcription Factor 4
DCFH-DA: 2’,7’-Dichlorodihydrofluorescein Diacetate
ER: Endoplasmic Reticulum
ERS: Endoplasmic Reticulum Stress
EVG: Elastica Van Gieson Staining
HAECs: Human Aortic Endothelial Cells
HASMCs: Human Aortic Smooth Muscle Cells
ICAM1: Intercellular Adhesion Molecule 1
IL-1β: Interleukin-1β
IL-6: Interleukin-6
IL-8: Interleukin-8
IL-32: Interleukin-32
MPs: Microplastics
MPS: PET-MPs signature
NAC: N-Acetylcysteine
Olink PEA: Olink Proximity Extension Assay
PE: Polyethylene
PET: Polyethylene Terephthalate
PET-MPs: Polyethylene Terephthalate Microplastics
PS: Polystyrene
PU: Polyurethane
PVC: Polyvinyl Chloride
PVDF: Polyvinylidene Fluoride
RNA: Ribonucleic Acid
ROS: Reactive Oxygen Species
SD rats: Sprague-Dawley rats
SDC1: Syndecan 1
TAA: Thoracic Aortic Aneurysm
TNF-α: Tumor Necrosis Factor Alpha
TUDCA: Tauroursodeoxycholic Acid
WGA: Wheat Germ Agglutinin
XBP1: X-Box Binding Protein 1.

