## Supplementary table 1: Detailed information on the antibodies, kits and other consumables used in this study. for "PET-Microplastics Trigger Endothelial Glycocalyx Loss via ER Stress and ROS Unleashing IL-1β-Driven SMC Switching and Early Aortic Structural Impairment"

|  | **Reagent name** | **Brand** | **Item number** | **Remarks** |
| --- | --- | --- | --- | --- |
| 1 | Syndecan 1 Recombinant Rabbit Monoclonal Antibody | HUABIO | ET1703-42-50 | 1:1000（WB）；1:100（IF） |
| 2 | ICAM-1/CD54 Rabbit Polyclonal Antibody | Proteintech | 30324-1-AP-50 | 1:1000（WB） |
| 3 | CD54/ICAM-1 Antibody | Cell Signaling | 4915S | 1:1000（WB） |
| 4 | NF-κB p65 Rabbit mAb | Selleck | F0006 | 1:1000（WB） |
| 5 | Phospho-NF-κB p65 (Ser536) Rabbit mAb | Selleck | F0155 | 1:1000（WB） |
| 6 | XBP1 Recombinant Rabbit Monoclonal Antibody | HUABIO | ET1703-23 | 1:1000（WB） |
| 7 | ATF4 Recombinant Rabbit Monoclonal Antibody | HUABIO | ET1612-37-50 | 1:1000（WB） |
| 8 | Calponin Recombinant Rabbit Monoclonal Antibody | HUABIO | ET1606-17-100 | 1:1000（WB）；1:100（IF） |
| 9 | Alpha smooth muscle Actin Recombinant Rabbit Monoclonal Antibody | HUABIO | ET1607-53-100 | 1:1000（WB）；1:100（IF） |
| 10 | a smooth muscle Actin/a-SMA Mouse mAb | Selleck | F2514 | 1:1000（WB）；1:100（IF） |
| 11 | GAPDH Rabbit pAb | ABclonal | AC001 | 1:2000（WB） |
| 12 | β-actin | Proteintech | 66009-1-IG-100 | 1:2000（WB） |
| 13 | Hoechst 33342 | Servicebio | G1127-1 | 1:1000（WB） |
| 14 | Anti-Active-β-Catenin (Anti-ABC) Antibody, clone 8E7 | Merck | 05-665 | 1:100（IF） |
| 15 | WHEAT GERM AGGLUTININ- TE | Thermo/Life/invitrogen | W849 | 1:100 |
| 16 | Lectin from Lycopersiconesculentum (tomato) | Sigma-Aldrich | L0401 | 1:100 |
| 17 | Anti-fluorescence quenching mounting solution | Beyotime | P0137-25 | 1:1 |
| 18 | Canakinumab | MedChemexpress | HY-108810 | 100ng/mL |
| 19 | Tauroursodeoxycholate | MedChemexpress | HY-19696 | 200μM |
| 20 | Acetylcysteine (N-acetylcysteine) | Selleck | S1623 | 5mM |
| 21 | Sulodexide | MedChemexpress | HY-100897 | 10LSU/mL |
| 22 | Copolymerization culture dish | Biosharp | BS-20A-GJM4 | None |
| 23 | Pentobarbital | Sigma-Aldrich | P3761 | None |
| 24 | Nitric acid | DaMao Chemical | 1107 | None |
| 25 | HASMC complete medium | ZhongQiaoXinZhou Biotechnology | ZMY016 | None |
| 26 | HAEC complete medium | ZhongQiaoXinZhou Biotechnology | ZMY001 | None |
| 27 | Triton | Servicebio | GC204003 | None |
| 28 | BSA | Servicebio | GC305010 | None |
| 29 | CCK-8 assay kit | Servicebio | G4103 | None |
| 30 | Reactive Oxygen Species Assay Kit | Servicebio | G1706 | None |
| 31 | Trizol | Invitrogen | 15596026CN | None |
| 32 | All-in-one RT SuperMix | Vazyme | R333-01 | None |
| 33 | ChamQ Universal SYBR qPCR Master Mix | Vazyme | Q411-02 | None |

Supplementary table 1: Detailed information on the antibodies, kits and other consumables used in this study.
