## Supplementary table 2:The basic information of 16 patients. for "PET-Microplastics Trigger Endothelial Glycocalyx Loss via ER Stress and ROS Unleashing IL-1β-Driven SMC Switching and Early Aortic Structural Impairment"

| **Number** | **Gender** | **Age** | **Disease** |
| --- | --- | --- | --- |
| ***Patient 1*** | Male | 74 | Aortic aneurysm;  Valvular heart disease:Moderate aortic insufficiency with calcification, mild mitral regurgitation,mild-moderate tricuspid regurgitation with pulmonary hypertension,atrial fibrillation; Coronary atherosclerotic heart disease;  Chronic cardiac insufficiency (NYHA Class II-III);  Hypertension, Stage 3, very high-risk group |
| ***Patient 2*** | Male | 58 | Aortic aneurysm;  Pulmonary infection; Atelectasis; Pleural effusion;  Hypertension, Stage 2, very high-risk group;  Left ureteral calculus & left renal calculus |
| ***Patient 3*** | Male | 68 | Aortic aneurysm; Gastrointestinal bleeding;  Coronary heart disease; Mild atherosclerosis;  Hypertension, Stage 3, very high-risk group;  Basal ganglia lacunar infarction;  Bilateral interstitial lung disease; Chronic cholecystitis;  Chronic hepatitis B; Allergic dermatitis |
| ***Patient 4*** | Male | 72 | Aortic aneurysm; Coronary heart disease;  Hypertension, Stage 3, very high-risk group;  Post-cerebral infarction sequela;  Atherosclerosis of carotid and subclavian arteries |
| ***Patient 5*** | Female | 56 | Aortic aneurysm; Hypertension, Stage 3, very high-risk group;  Coronary heart disease; Atelectasis |
| ***Patient 6*** | Female | 59 | Aortic aneurysm; Bicuspid aortic valve;  Pulmonary infection; Hyperbilirubinemia;  Hypertension, Stage 2, very high-risk group;  Electrolyte imbalance |
| ***Patient 7*** | Male | 63 | Aortic aneurysm; Hypertension, Stage 3, very high-risk group;  Renal insufficiency; Hypoalbuminemia |
| ***Patient 8*** | Male | 46 | Aortic aneurysm; Valvular heart disease:  Mild aortic regurgitation;  Klebsiella pneumoniae pneumonia |
| ***Patient 9*** | Male | 50 | Aortic aneurysm; Gastrointestinal bleeding;  Hypertension, Stage 3, very high-risk group;  Sleep disorder |
| ***Patient 10*** | Male | 70 | Aortic aneurysm; Hypertension, Stage 3, very high-risk;  Coagulopathy; Hypernatremia; Hypoalbuminemia;  Coronary heart disease |
| ***Patient 11*** | Male | 53 | Aortic aneurysm; Hypertension, Stage 3, very high-risk group;  Pulmonary infection; Atelectasis |
| ***Patient 12*** | Female | 63 | Aortic aneurysm; Pleural effusion; Pulmonary infection;  Thrombocytopenia with purpura; Gastrointestinal bleeding;  Post-bioprosthetic aortic valve replacement;  Suspected spinal cord ischemia? Incomplete quadriplegia; Asthma |
| ***Patient 13*** | Female | 66 | Aortic aneurysm; Bilateral pleural effusion;  Left renal ischemia; Hypoxic-ischemic encephalopathy;  Hypertension, Stage 3, very high-risk group |
| ***Patient 14*** | Male | 65 | Aortic aneurysm; Hypertension, Stage 3, very high-risk group;  Severe stenosis of proximal left common carotid artery with mild atherosclerosis;  Pleural effusion; Gastrointestinal bleeding;  Postoperative left upper lung squamous cell carcinoma (pT4N0M0, Stage IIIA);  Left lower lung tuberculosis; Multiple lacunar infarcts with cerebral atrophy |
| ***Patient 15*** | Male | 48 | Aortic aneurysm; Pleural effusion; Pulmonary infection;  Hypertension, Stage 3, very high-risk group; Hepatic insufficiency |
| ***Patient 16*** | Female | 63 | Aortic aneurysm; Coronary heart disease;  Valvular disease: Moderate aortic regurgitation, mild tricuspid regurgitation,mild mitral regurgitation |

Supplementary table 2:The basic information of 16 patients.
