## Supplementary table 3: Primer sequences used in this study. for "PET-Microplastics Trigger Endothelial Glycocalyx Loss via ER Stress and ROS Unleashing IL-1β-Driven SMC Switching and Early Aortic Structural Impairment"

|  | **Primer** | **Sequence（5'-3'）** |
| --- | --- | --- |
| 1 | H-ACTA2-F | AAAAGACAGCTACGTGGGTGA |
| 2 | H-ACTA2-R | GCCATGTTCTATCGGGTACTTC |
| 3 | H-Tagln-F | AGTGCAGTCCAAAATCGAGAAG |
| 4 | H-Tagln-R | CTTGCTCAGAATCACGCCAT |
| 5 | H-TNC-F | TCCCAGTGTTCGGTGGATCT |
| 6 | H-TNC-R | TTGATGCGATGTGTGAAGACA |
| 7 | H-FN-F | CGGTGGCTGTCAGTCAAAG |
| 8 | H-FN-R | AAACCTCGGCTTCCTCCATAA |
| 9 | H-COL1a1-F | GAGGGCCAAGACGAAGACATC |
| 10 | H-COL1a1-R | CAGATCACGTCATCGCACAAC |
| 11 | H-ATF4-F | ATGACCGAAATGAGCTTCCTG |
| 12 | H-ATF4-R | GCTGGAGAACCCATGAGGT |
| 13 | H-XBP1s-F | CCCTCCAGAACATCTCCCCAT |
| 14 | H-XBP1s-R | ACATGACTGGGTCCAAGTTGT |
| 15 | H-IL1β-F | AACCTCTTCGAGGCACAAGG |
| 16 | H-IL1β-R | GTCCTGGAAGGAGCACTTCAT |
| 17 | H-IL6-F | AGACAGCCACTCACCTCTTCAG |
| 18 | H-IL6-R | TTCTGCCAGTGCCTCTTTGCTG |
| 19 | H-IL8-F | GAGAGTGATTGAGAGTGGACCAC |
| 20 | H-IL8-R | CACAACCCTCTGCACCCAGTTT |
| 21 | H-EXT1-F | GCTCTTGTCTCGCCCTTTTGT |
| 22 | H-EXT1-R | GTGGTGCAAGCCATTCCTAC |
| 23 | H-EXT2-F | ATGTGTGCGTCGGTCAAGTAT |
| 24 | H-EXT2-R | AGAATGGGGCCAAAACTGAAA |
| 25 | H-HAS2-F | CTCTTTTGGACTGTATGGTGCC |
| 26 | H-HAS2-R | AGGGTAGGTTAGCCTTTTCACA |
| 27 | H-HAS3-F | CAGCCTATGTGACGGGCTAC |
| 28 | H-HAS3-R | CCTCCTGGTATGCGGCAAT |

Supplementary table 3: Primer sequences used in this study.
