## Supplementary table4: Calculate the gene set required for the endocalyx. for "PET-Microplastics Trigger Endothelial Glycocalyx Loss via ER Stress and ROS Unleashing IL-1β-Driven SMC Switching and Early Aortic Structural Impairment"

| **SYMBOL** | **Term** |
| --- | --- |
| Has1 | Glycocalyx score |
| Has2 | Glycocalyx score |
| Has3 | Glycocalyx score |
| Ugp2 | Glycocalyx score |
| Ugdh | Glycocalyx score |
| Gfpt1 | Glycocalyx score |
| Hs2St1 | Glycocalyx score |
| Hs3St1 | Glycocalyx score |
| Hs6St1 | Glycocalyx score |
| Glce | Glycocalyx score |
| Ndst1 | Glycocalyx score |
| Ndst2 | Glycocalyx score |
| Ext1 | Glycocalyx score |
| Ext2 | Glycocalyx score |
| Chsy1 | Glycocalyx score |
| Chsy3 | Glycocalyx score |
| Sulf1 | Glycocalyx score |
| Sulf2 | Glycocalyx score |
| Hpse | Glycocalyx score |
| Sdc1 | Glycocalyx score |
| Sdc2 | Glycocalyx score |
| Sdc3 | Glycocalyx score |
| Sdc4 | Glycocalyx score |
| Gpc1 | Glycocalyx score |
| Hspg2 | Glycocalyx score |
| Bgn | Glycocalyx score |
| Vcan | Glycocalyx score |
| Dcn | Glycocalyx score |
| Hyal2 | Glycocalyx score |
| Cd44 | Glycocalyx score |
| Habp2 | Glycocalyx score |
| Habp4 | Glycocalyx score |
| Tnfaip6 | Glycocalyx score |
| Ambp | Glycocalyx score |
| Has1 | Glycocalyx biosynthesis |
| Has2 | Glycocalyx biosynthesis |
| Has3 | Glycocalyx biosynthesis |
| Ugp2 | Glycocalyx biosynthesis |
| Ugdh | Glycocalyx biosynthesis |
| Gfpt1 | Glycocalyx biosynthesis |
| Hs2St1 | Glycocalyx biosynthesis |
| Hs3St1 | Glycocalyx biosynthesis |
| Hs6St1 | Glycocalyx biosynthesis |
| Glce | Glycocalyx biosynthesis |
| Ndst1 | Glycocalyx biosynthesis |
| Ndst2 | Glycocalyx biosynthesis |
| Ext1 | Glycocalyx biosynthesis |
| Ext2 | Glycocalyx biosynthesis |
| Chsy1 | Glycocalyx biosynthesis |
| Chsy3 | Glycocalyx biosynthesis |
| Sulf1 | Glycocalyx modification |
| Sulf2 | Glycocalyx modification |
| Hpse | Glycocalyx modification |
| Sdc1 | Luminal glycocalyx |
| Sdc2 | Luminal glycocalyx |
| Sdc3 | Luminal glycocalyx |
| Sdc4 | Luminal glycocalyx |
| Gpc1 | Luminal glycocalyx |
| Hspg2 | Luminal glycocalyx |
| Bgn | Abluminal glycocalyx |
| Vcan | Abluminal glycocalyx |
| Dcn | Abluminal glycocalyx |
| Hyal2 | Glycocalyx stabilisation |
| Cd44 | Glycocalyx stabilisation |
| Habp2 | Glycocalyx stabilisation |
| Habp4 | Glycocalyx stabilisation |
| Tnfaip6 | Glycocalyx stabilisation |
| Ambp | Glycocalyx stabilisation |

Supplementary table4: Calculate the gene set required for the endocalyx.
