## Supplementary table 7:Py-GC/MS calibration curves of polymers. for "PET-Microplastics Trigger Endothelial Glycocalyx Loss via ER Stress and ROS Unleashing IL-1β-Driven SMC Switching and Early Aortic Structural Impairment"

| **Supplementary Table 7.Py-GC/MS calibration curves of polymers** | | | | | | |
| --- | --- | --- | --- | --- | --- | --- |
| **Polymer** | **Characteristic compound** | **Retention time(min)** | **Calibration curve equation** | **R2** | **LOD(μg)** | **LOQ(μg)** |
| PET | benzoic acd | 9.954 | Y=43653 * x + 19734.8 | 0.996004 | 0.02 | 0.06 |
| PE | n-monoene | 14.398 | Y=90663.9x + -422837 | 0.997293 | 0.5 | 0.15 |
| PVC | naphthalene | 3.938 | Y=187227 * x + -966404 | 0.996353 | 0.02 | 0.06 |
| PA-66 | cyclopentanone | 9.954 | Y=25151.9 * x + -13826.2 | 0.994654 | 0.02 | 0.06 |

Abbreviations: LOD, limit of detection: LOQ, limit of quantitation; PA, polyamide;

PE, polyethylene; PET,polyethylene terephthalate; PVC, polyvinyl chloride: Py-GC/Ms, pyrolysis gas chromatography/mass spectrometry.
