## Supplementary figures and images for "PET-Microplastics Trigger Endothelial Glycocalyx Loss via ER Stress and ROS Unleashing IL-1β-Driven SMC Switching and Early Aortic Structural Impairment"

### Supplementary Figure l. Gross morphology of arterial samples

Supplementary Figure I. Gross morphology of arterial samples

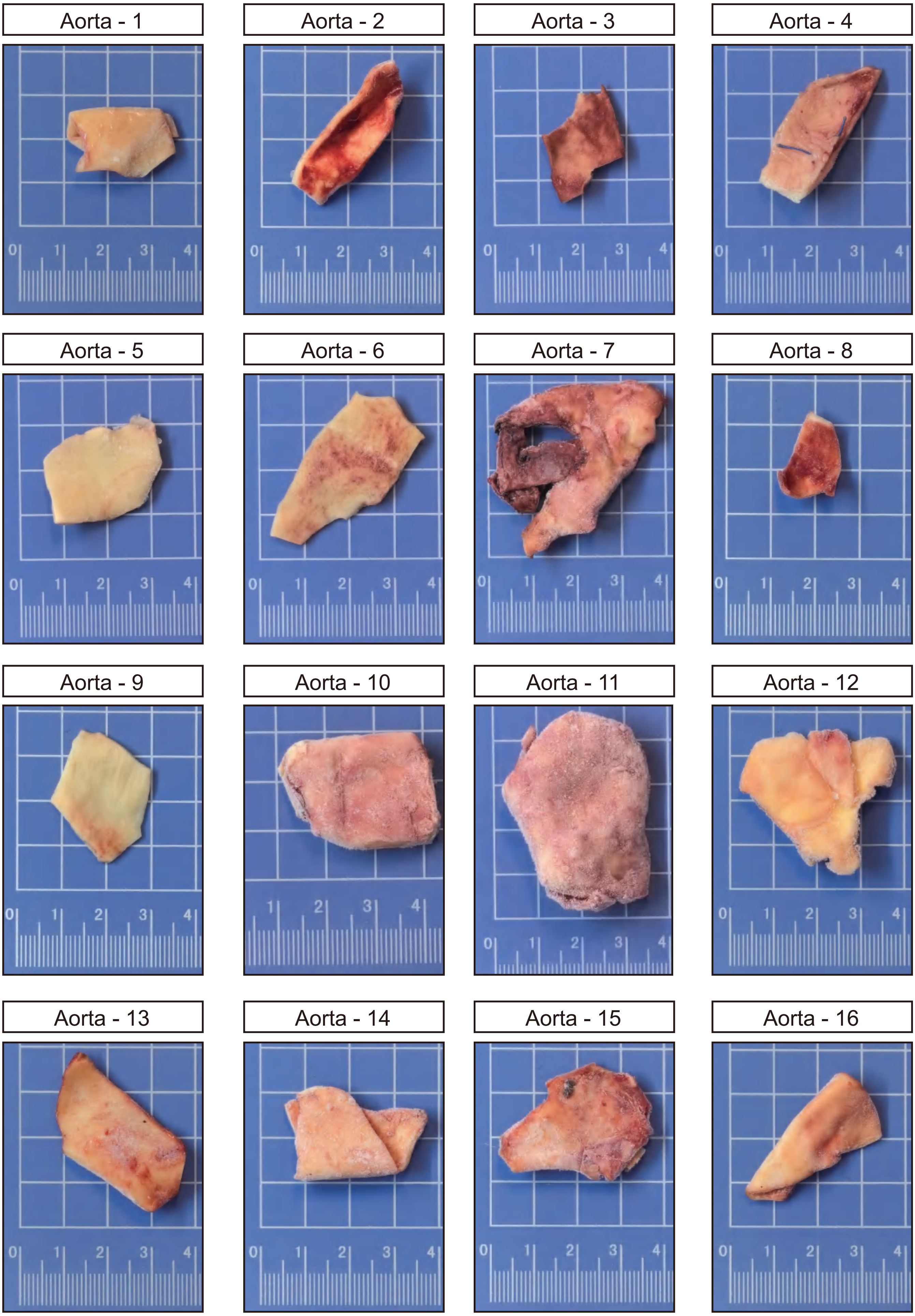
