## Supplementary Figure 2. for "PET-Microplastics Trigger Endothelial Glycocalyx Loss via ER Stress and ROS Unleashing IL-1β-Driven SMC Switching and Early Aortic Structural Impairment"

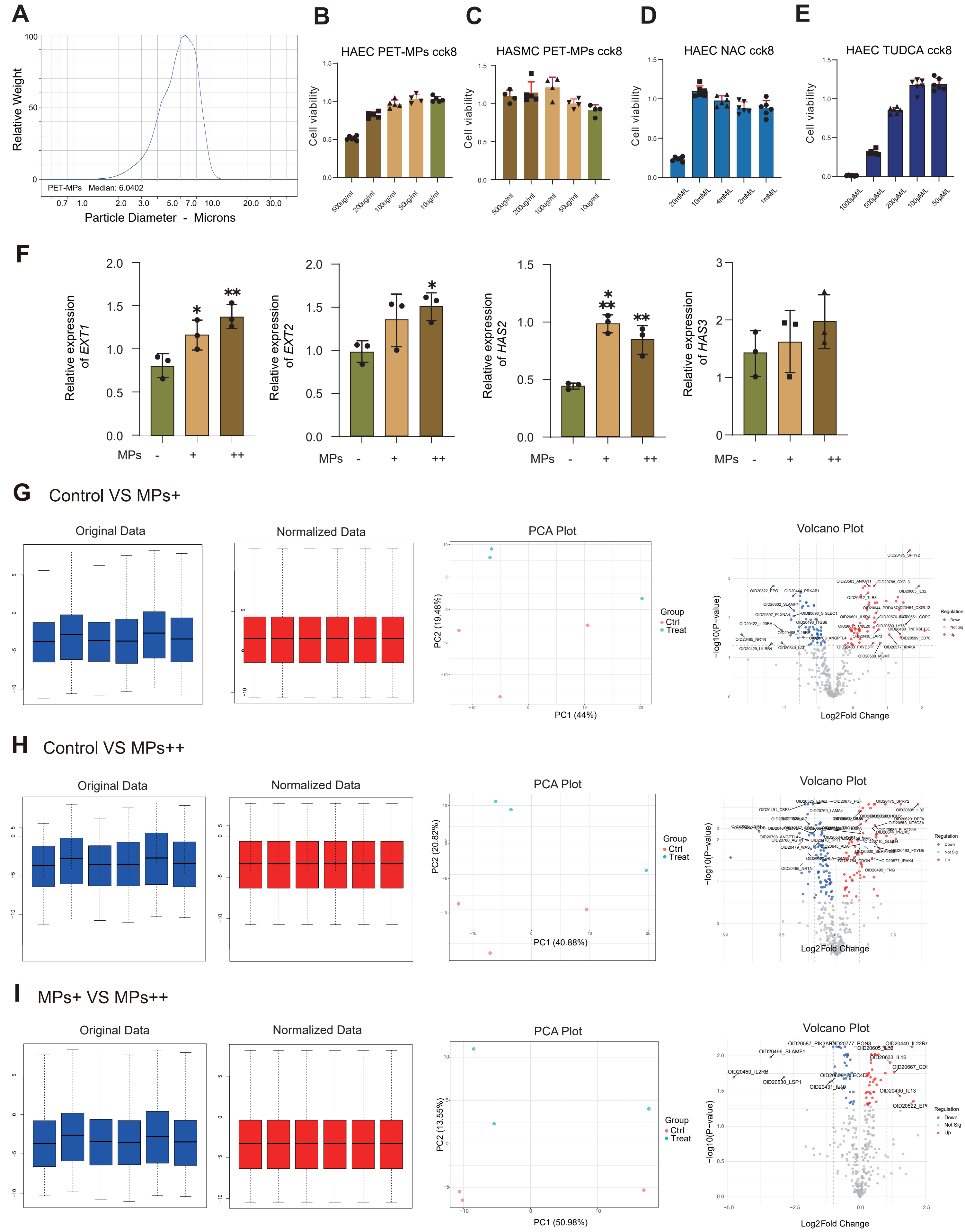


(A) Particle size distribution profile of PET-MPs used in this study.
(B) CCK-8 viability assay of HAECs exposed to PET-MPs.
(C) CCK-8 viability assay of HASMCs exposed to PET-MPs.
(D) CCK-8 viability assay of HAECs treated with NAC.
(E) CCK-8 viability assay of HAECs treated with TUDCA.
(F) qPCR analysis of endothelial glycocalyx synthesis-related genes in HAECs following PET-MPs exposure.
(G-I) Olink panel analysis (proximity extension assay).
